# How to quantitatively measure, assess and correct the fluorescence crosstalk in the wide-field microscopy

**DOI:** 10.1101/2020.05.20.105627

**Authors:** Hieng Chiong Tie, Lei Lu

## Abstract

During the multi-color fluorescence imaging, a fluorophore can be detected in the non-corresponding channel due to the broad spectra of the fluorophore and filters, resulting in the artefact called the crosstalk (crossover or bleed-through). Unfortunately, the fluorescence crosstalk is ubiquitous for the conventional fluorescence microscopy. Significant crosstalk can distort the image and lead to incorrect interpretation. Here, we introduce a simple quantitative metric, the crosstalk factor, to measure the crosstalk effect of a fluorophore on a channel in the wide-field fluorescence microscopy. We describe a cell biologist friendly protocol which should be easily implemented by cell biologists using commonly available image processing software such as ImageJ.

## Introduction

The multi-color fluorescence imaging is widely used in the biology to simultaneously reveal the spatial localization of different target molecules at the cellular or subcellular level. In this imaging technique, the signal from each fluorophore is recorded via the corresponding imaging channel. For a typical wide-field microscopic system, a channel is defined by the combination of the excitation filter, dichroic mirror and emission filter. Ideally, a channel should collect as many photons from its target fluorophore as possible while at the same time rejects those from non-target one, therefore ensuring that the image acquired by each channel strictly maps its target fluorophore.

However, in a real microscopic system, a channel might also register the emission signal from the non-target fluorophore. This undesirable signal is referred to as the crosstalk (crossover or bleed-through) and it is produced primarily due to the broad excitation and emission spectra of the non-target fluorophore and/or channel filter (Murphy et al.). When fluorophores and microscopic filters are selected, the imbalanced fluorophore labeling of the sample can aggravate the crosstalk and the imaging channel with a weak signal is usually more prone to the crosstalk. Unfortunately, the fluorescence crosstalk of different degrees of severity is ubiquitous for the conventional multicolor microscopic imaging. While the crosstalk below certain threshold can be tolerated or neglected, a severe crosstalk can often complicate or mislead the quantitative and qualitative interpretation of the fluorescence imaging. For example, the crosstalk can result in the false-positive subcellular colocalization of two proteins. When using the sensitized emission method to study FRET (fluorescence resonance energy transfer), the weak FRET signal can often be obscured by the crosstalk.

There are commonly recommended practices to reduce the crosstalk. For instance, it can be significantly reduced by selecting fluorophores whose excitation and emission spectra match those of the corresponding channels but minimally overlap those of non-corresponding ones. Alternatively, optimizing the filters of imaging channels, such as the adoption of excitation and emission filters with narrower bandwidths, can very effectively alleviate the crosstalk, although the signal strength might be sacrificed. However, once the fluorophore and microscopic hardware configuration are fixed, the post-acquisition method to assess and correct the crosstalk remains the only choice (Murphy et al.).

Crosstalk correction methods such as spectral linear unmixing are available in the literature. But these methods seem not popular in cellular imaging partially because the concepts and software tools are beyond most cell biologists. Here, we describe a cell biologist friendly crosstalk correction protocol based on the spectral linear unmixing. We introduce a simple quantitative metric, the crosstalk factor, to measure the crosstalk effect of a fluorophore on a channel in the wide-field fluorescence microscopy.

## Results and discussion

### The theory behind the crosstalk factor

Assuming that the fluorophore x and y are to be imaged by a conventional wide-field microscope through two corresponding channels X and Y respectively. Each channel has fixed optical components along the optical train, including the channel specific filters/dichroic mirrors and the shared excitation light source, apertures, the tube lens, the objective and the charge-coupled device (CCD) or complementary metal-oxide semiconductor (CMOS) camera. When a specimen is singly labeled with the fluorophore x, two images can be sequentially acquired via channel X with the exposure time *t_X_*, followed by channel Y with the exposure time *t_Y_*. The pair of images are subsequently background-subtracted. In the channel X image, the ROI of an object has the mean intensity *I_X_*. The channel Y image should be the crosstalk image since it is solely contributed by the non-target fluorophore x. In the crosstalk image, the same ROI has the mean intensity *I_Y_*. Due to fixed optical train components, the crosstalk factor *C_x,Y_* can be defined by the below equation (1) and should be a constant.

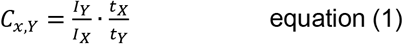

In the notion of the crosstalk factor *C_x,Y_*, the direction of the crosstalk is indicated by the subscript, of which the first part denotes the fluorophore which is the source of the crosstalk, while the second part indicates the non-corresponding channel. For a wide-field microscope, the crosstalk factor is dependent on the spectra of the fluorophore and channel filters but independent on the image exposure time and the fluorophore labeling density. Crosstalk factors of commonly used fluorophores to different channels can therefore be measured in advance and employed later to quantitatively assess and correct the crosstalk.

### Measuring the crosstalk factor

A specimen singly labeled with fluorophore x is employed to measure the crosstalk factor *C_x,Y_*. Since the crosstalk signal is usually weak, the labeling of fluorophore x in the specimen must be sufficiently bright and the exposure time *t_Y_* must be sufficiently long to acquire an analyzable crosstalk image in the channel Y. An example is described below to illustrate how we measured the crosstalk factor. We found that HeLa cells singly overexpressing the chromatin marker, H2B tagged with either the fluorescence protein (GFP or mCherry) or Myc-tag, provide a reliable fluorescence source for the measurement of crosstalk factors. During the immunofluorescence labeling, Myc-tag can be stained with the testing fluorophore such as Alexa Fluor dyes. In Figure 1A, a cell expressing H2B-mCherry was imaged in the channel R (red channel) and B (far red channel) with exposure time *t_R_* and *t_B_* to produce *Image_R_* and *Image_B_* respectively. We use *Image_mCherry,R_* to represent the crosstalk-free true mCherry image in the channel R. In this case, it is the same as *ImageR* since the sample is singly labeled with mCherry. *Image_B_* is the crosstalk image, which is referred to as *CImage_mCherry,B_* (C denotes crosstalk). The mean intensity of the ROI within the nucleus was measured to be *I_R_* and *I_B_* respectively. Hence, the crosstalk factor, *C_mCherry,B_*, can be calculated according to equation (1).

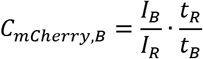

**Figure 1.**
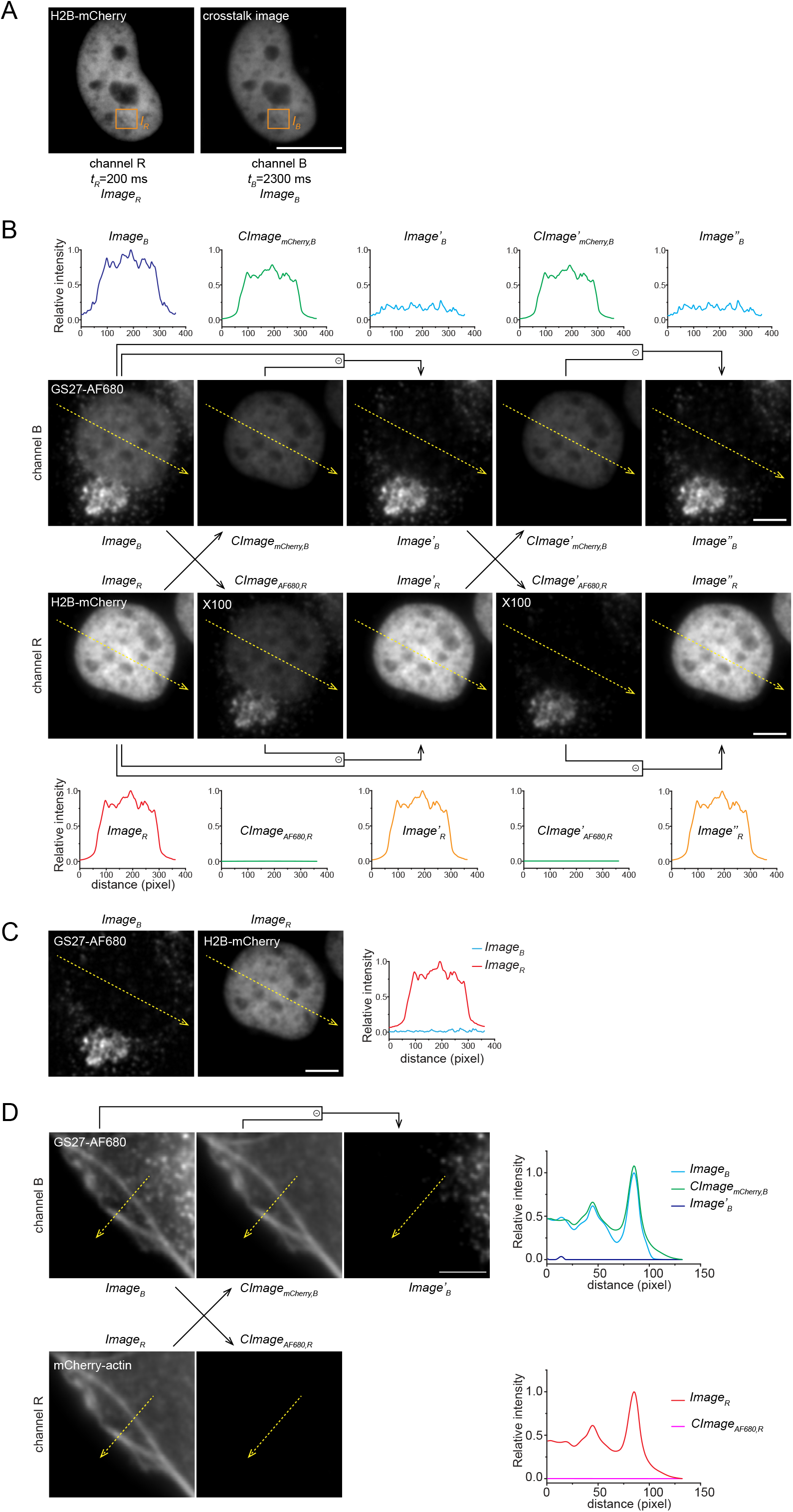
Measuring the crosstalk factor and using it to assess and correct the crosstalk. HeLa cells are used. (**A**) An example demonstrating the measurement of the crosstalk factor, *C_mCherry,B_*. Cells transiently expressing H2B-mCherry were fixed and imaged via channel R (H2B-mCherry image) and B (crosstalk image) with exposure time *t_R_* and *t_B_* respectively. A square ROI is drawn within the nucleus. *I_R_* and *I_B_* are the mean intensity within the ROI in channel R and B respectively. The equation to calculate *C_mCherry,B_* are described in the text. Scale bar, 10 μm. (**B**) Crosstalk assessment and correction in 2-color imaging. Cells expressing H2B-mCherry were subjected to AF680-conjugated immuno-staining of endogenous GS27. Images were acquired by channels using filter cubes. Serious crosstalk from mCherry to channel B can be observed in *Image_B_*. Within each row, intensities of images are adjusted to the same range, except for those of *CImage_AF680,R_* and *CImage’_AF680,R_*, which are amplified 100-fold to reveal the very weak crosstalk signals. Lines immediately above or below images indicate the subtraction of two input images to result in the calculated one. Plots immediately above or below images show the line intensity profiles of dotted yellow arrow lines. Scale bar, 5 μm. (**C**) Crosstalk-free images. The same field of view of (B) was imaged by channels using the filter wheel, crosstalk factors of which are 1000-fold less than those of corresponding channels using filter cubes. Line intensity profiles are plotted at the right. Scale bar, 5 μm. (**D**) Cells expressing mCherry-actin were subjected to AF680-conjugated immuno-staining of endogenous GS27. Images were acquired by channels using filter cubes. The organization of images is similar to (B). The crosstalk correction is only needed in channel B as the intensity of *CImage_AF680,R_* is negligible in comparison to *Image_R_*. Scale bar, 2 μm.

Similarly, we can systematically measure crosstalk factors of commonly used fluorophores in channels of different filter configurations (Table 1).

**Table 1.**
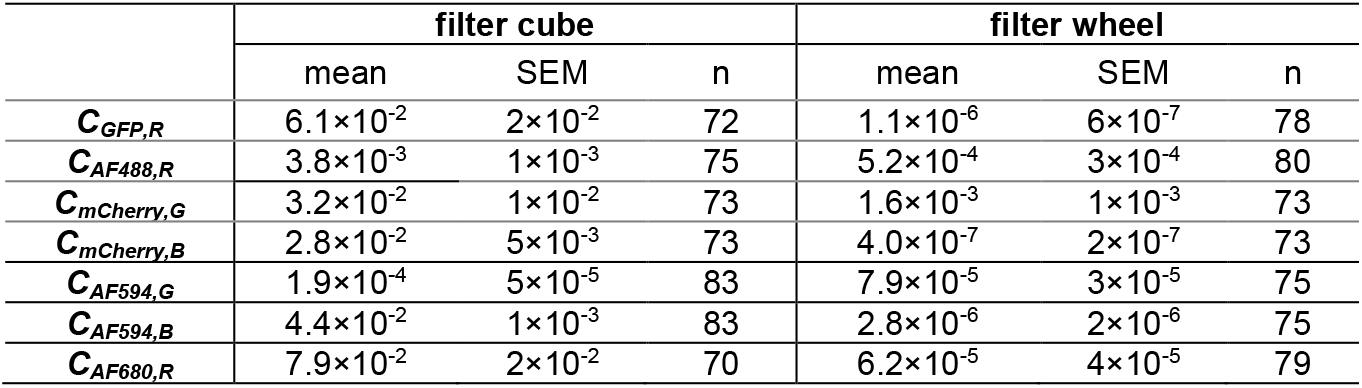
Crosstalk factors of fluorophores for channels using filter cubes and the filter wheel. SEM, standard error of mean; n, the number of nuclei quantified.

|  | filter cube |  |  | filter wheel |  |  |
| --- | --- | --- | --- | --- | --- | --- |
|  | mean | SEM | n | mean | SEM | n |
| $C_{GFP,R}$ | $6.1 \times 10^{-2}$ | $2 \times 10^{-2}$ | 72 | $1.1 \times 10^{-6}$ | $6 \times 10^{-7}$ | 78 |
| $C_{AF488,R}$ | $3.8 \times 10^{-3}$ | $1 \times 10^{-3}$ | 75 | $5.2 \times 10^{-4}$ | $3 \times 10^{-4}$ | 80 |
| $C_{mCherry,G}$ | $3.2 \times 10^{-2}$ | $1 \times 10^{-2}$ | 73 | $1.6 \times 10^{-3}$ | $1 \times 10^{-3}$ | 73 |
| $C_{mCherry,B}$ | $2.8 \times 10^{-2}$ | $5 \times 10^{-3}$ | 73 | $4.0 \times 10^{-7}$ | $2 \times 10^{-7}$ | 73 |
| $C_{AF594,G}$ | $1.9 \times 10^{-4}$ | $5 \times 10^{-5}$ | 83 | $7.9 \times 10^{-5}$ | $3 \times 10^{-5}$ | 75 |
| $C_{AF594,B}$ | $4.4 \times 10^{-2}$ | $1 \times 10^{-3}$ | 83 | $2.8 \times 10^{-6}$ | $2 \times 10^{-6}$ | 75 |
| $C_{AF680,R}$ | $7.9 \times 10^{-2}$ | $2 \times 10^{-2}$ | 70 | $6.2 \times 10^{-5}$ | $4 \times 10^{-5}$ | 79 |

### Quantitative assessment and correction of the crosstalk

Crosstalk factors can be used to evaluate the potential crosstalk artefact of a microscope filter setup. For example, Table 1 indicates that, for our microscope, crosstalk factors of channels using the filter wheel are at least 10-fold less than those using filter cubes. Therefore, images acquired using the filter wheel should produce much less crosstalk than those using filter cubes (see below for an example). If a crosstalk factor is significantly large, e.g. ≥ 0.10, we recommend that it is better to select an alternative fluorophore or filter configuration than to perform the post-acquisition processing described below.

Here is an example showing how crosstalk factors can be employed to assess and correct the crosstalk in the multi-color fluorescence imaging. HeLa cells transiently expressing H2B-mCherry were immuno-stained for GS27 (a Golgi marker) using Alexa Fluor 680 (hereafter AF680) (Fig. 1B). Using filter cubes, fluorescence images of mCherry and AF680 were acquired through channel R (*Image_R_*) and B (*Image_B_*) with exposure time *t_R_* and *t_B_* respectively. There are two possible crosstalks: from mCherry to the channel B and from AF680 to the channel R. *Image_B_* can be considered as the addition of the true AF680 image, *Image_AF680,B_*, and the crosstalk image from mCherry to the channel B, *CImage_mCherry,B_*. *CImage_mCherry,B_* can be derived from *Image_R_*, which approximates *Image_mCherry,R_*, using the below transformation of equation (1).

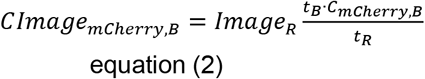

This calculation can be easily conducted using the image arithmetic tool in ImageJ (Process>Math>Multiply). The resulting crosstalk image, *CImage_mCherry,B_*, is subsequently compared with *Image_B_* for pixel values or brightness (under identical intensity scaling). The crosstalk can be ignored if the intensity of *CImage_mCherry,B_* is significantly less than that of *Image_B_* for all pixels within a ROI. In our case (Fig. 1B), the nuclear signal in *Image_B_* is clearly an artefact since its intensity is similar to *CImage_mCherry,B_*. The most straightforward solution to avoid this artefact is to select the filter wheel, which has a much smaller crosstalk factor *C_mCherry,B_* (4.0×10^−7^, Table 1) than the filter cube (2.8×10^-2^, Table 1). Indeed, using the filter wheel, the GS27-AF680 image of the same specimen field does not have detectable crosstalk from mCherry (Fig. 1C). The absence of GS27 in the nucleus is consistent with the report that, GS27, a soluble N-ethylmaleimide sensitive factor attachment protein receptor (SNARE), does not localize to the nucleus (Lowe et al., 1997).

Through the post-acquisition image processing, the crosstalk can be corrected simply by subtracting *CImage_mCherry,B_* from *Image_B_* to generate the crosstalk-corrected image *Image’_B_* (Fig. 1B) as below.

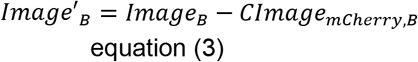

Likewise, *CImage_AF680,R_* and the crosstalk-corrected image, *Image’_R_* (Fig. 1B), can be calculated as below.

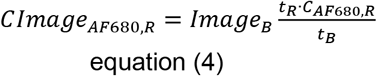

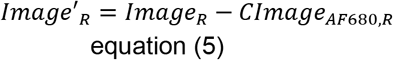

Note that *CImage_AF680,R_* is very weak in comparison to *Image_R_*, hence the crosstalk is negligible (Fig. 1B). To make *CImage_AF680,R_* visible, its intensity displayed in Figure 1B is amplified 100-fold.

The crosstalk correction is assessed by plotting line intensity profiles (Fig. 1B). The significantly reduced nuclear signal in the line intensity profile of *Image’_B_* demonstrates the crosstalk correction. An iterative calculation process may be adopted by substituting *Image_R_* and *Image_B_* with *Image’_R_* and *Image’_B_* in equations (2-3) and (4-5) respectively for better approximations of *Image_mCherry,R_*, and *Image_AF680,B_*. However, the improvement yielded by such iterative process is very small as judged by comparing *Image_B_*, *Image’_B_* and *Image’’_B_* (Fig. 1B). Therefore, the first round of calculation should usually be sufficient.

In most cases, the crosstalk correction is only required in one direction, which further simplifies the calculation. For example, in Figure 1B, only *CImage_mCherry,B_* is significant while *CImage_AF680,R_* is negligible. Therefore, only *ImageB* needs the crosstalk correction. As another example, the crosstalk correction in a set of GS27 and mCherry-actin fluorescence images are demonstrated in Figure 1D.

## Materials and methods

### DNA plasmids and antibodies

For our microscope, the crosstalk correction in images acquired using filter cubes can be evaluated by comparing processed images with those acquired using the filter wheel, which are assumed to be crosstalk-free due to their much smaller crosstalk factors (Table 1). Comparing to Figure 1C, there is still residual amount of crosstalk signal in the nuclear region of *Image’_B_* (or *Image’’_B_*) (Fig. 1B), indicating that the crosstalk correction is still incomplete (under-corrected). On the other hand in Figure 1D, the intensity of *CImage_mCherry,B_* is even slightly higher than *Image_B_* as demonstrated from the line intensity profile, therefore leading to zero intensity for *Image’_B_* intensity profile (over-corrected). The unsatisfactory crosstalk correction is mainly due to the difficulty in determining an accurate crosstalk image. The calculated crosstalk image can be significantly off due to multiple practical issues, such as the photo-bleaching of the crosstalk source fluorophores during the image acquisition, the imprecise measurement of crosstalk factors, the bias in the background subtraction and the noises introduced during the imaging. These issues can limit the quantitative application of our correction method.

In summary, we describe here a simple method to empirically measure, assess and correct the crosstalk of the multi-channel imaging in the wide-field microscopy. The crosstalk factor indicates the potential crosstalk of the instrumentation setup for the specific fluorophore. It can be used to guide the selection of the fluorophore and microscopic filters to minimize the crosstalk artefact. After the image acquisition, the crosstalk image can be generated to assess the acquired image for the crosstalk and, in some cases, satisfactorily correct the crosstalk artefact. However, for images with a serious crosstalk, we still recommend to first optimize the selection of filters and fluorophores or improve the labeling of the sample.

DNA plasmid pBOS-H2B-GFP that expresses H2B-GFP was from BD Pharmingen. mCherry-actin (Lu et al., 2009) and H2B-mCherry (Lu et al., 2011) were previously described. To construct H2B-Myc, the coding sequence of H2B was first PCR-amplified from pBOS-H2B-GFP using a pair of oligonucleotide primers (5’-GAT GCA CTC GAG TCG CCA CCA TGC CAG AGC CAG CGA AG-3’ and 5’-CAT GTT GTC GAC CTT AGC GCT GGT GTA CTT G-3’). The PCR product was then digested by XhoI/SalI and ligated into pDMyc-neo-N1 vector (Tie et al., 2018) using the same restriction sites. Mouse anti-human GS27 monoclonal antibody (#611034) was from BD Biosciences. Mouse anti-Myc tag monoclonal antibody (#sc-40) was from Santa Cruz. Alexa Fluor 488, 594, 647 and 680-conjugated goat antibodies against mouse IgG were from Thermo Fisher Scientific.

### Immunofluorescence labeling

HeLa cells were seeded onto Φ 12 mm glass coverslips and cultured in DMEM supplemented with 10% fetal bovine serum at 37 °C in a CO2 incubator. Transfection was performed using Lipofectamine2000 (Invitrogen). Cells expressing H2B-GFP or –mCherry were fixed by 4% paraformaldehyde in PBS at room temperature for 20 min. Residual paraformaldehyde was neutralized using 100 mM ammonium chloride. Cells expressing H2B-Myc were fixed by methanol at – 20 °C and subsequently washed by PBS. For immunofluorescence labeling, both paraformaldehyde and methanol fixed cells are incubated with primary antibody diluted in the antibody dilution buffer (PBS supplemented with 5% FBS, 2% bovine serum albumin and 0.1% saponin (Sigma-Aldrich)). After PBS washing, cells were incubated with fluorescence-conjugated secondary antibody diluted in the antibody dilution buffer. At last, the coverslip was washed in PBS and mounted in Mowiol 4-88 (EMD Millipore).

### Imaging

Images were acquired under an inverted wide-field microscope system, comprising Olympus IX83 equipped with 100x (U plan super apochromat, NA 1.40) oil objective lens, a motorized stage, motorized filter cubes, motorized filter wheels housing different excitation and emission filters, a scientific complementary metal-oxide semiconductor camera (Neo; Andor) and a 200 W metal-halide excitation light source (Lumen Pro 200; Prior Scientific). All filters were from Chroma Technology Corp. The configuration of filter cubes is listed below (catalog numbers are in the order of the excitation filter, dichroic mirror and emission filter): channel G (ET470/40x, T495lpxr and ET525/50m), channel R (ET560/40x, T585lpxr and ET630/75m) and channel B (ET620/60x, T660lpxr and ET700/75m). The same dichroic mirror (89100bs) is shared among all channels using the filter wheel, the configuration of which is listed below (catalog numbers are in the order of the excitation and emission filter): channel G (ET490/20x and ET525/36m), channel R (ET555/25x and ET605/52m) and channel B (ET645/30x and ET705/72m). The microscope system was controlled by MetaMorph software (Molecular Devices) and only center quadrant of the camera sensor was used for imaging.

### Image processing and analysis

All image analysis and manipulations were performed in ImageJ (http://imagej.nih.gov/ij/). Images were Gaussian filtered (radius equals 2 pixels) and background-subtracted before further processing and analysis.

## Author contributions

LL conceived and supervised the study and designed experiments. HCT performed experiments. LL and HCT analyzed the data and prepared figures. LL wrote the manuscript.

## Acknowledgements

This work was supported by the following grants to L.L.: MOE AcRF Tier1 RG35/17, Tier2 MOE2015-T2-2-073 and MOE2018-T2-2-026.

